# *Ex vivo* recapitulation of intramuscular mRNA vaccination with naïve and recall antigens using a human Lymphoid Follicle Chip platform

**DOI:** 10.1101/2025.04.16.649172

**Authors:** Yunhao Zhai, Min Wen Ku, Ka Yang, Aditya Patil, Pranav Prabhala, Liqun He, Yuncheng Man, Sven B. Spörri, Supriya Gharpure, Abdul Rahman Isaacs, Shay Ferdosi, Andrzej S. Pitek, Giulietta Maruggi, Sylvie Bertholet, Jessica Firestone, Kambiz Mousavi, Eric Miller, Kate Luisi, Caleb A. Hellman, LaTonya D. Williams, Georgia D. Tomaras, Peng Yin, Steven P. Gygi, Josie McAuliffe, Donald E. Ingber, Girija Goyal

**Affiliations:** Wyss Institute for Biologically Inspired Engineering, Harvard University, Boston, MA 02215, USA; Department of Cell Biology, Harvard Medical School, Boston, MA 02115, USA; Vascular Biology Program and Department of Surgery, Boston Children’s Hospital and Harvard Medical School, Boston, MA 02115, USA; GSK, Rockville Center for Vaccine Research, Rockville, MD 20850, USA; Center for Human Systems Immunology, Departments of Surgery and Integrative Immunobiology, Duke University School of Medicine, Durham, North Carolina 27701, USA; Department of Systems Biology, Harvard Medical School, Boston, MA, USA; Harvard John A. Paulson School of Engineering and Applied Sciences, Cambridge, MA 02139, USA

## Abstract

Predicting the efficacy and toxicity of intramuscular mRNA vaccines remains challenging. Here, we describe an *ex vivo* human cell-based model that replicates immune responses to lipid nanoparticle (LNP)-based mRNA vaccines that require intramuscular injection. Vaccines are administered into a biomimetic muscle module containing human skeletal myoblasts and antigen-presenting cells (APCs) to mimic intramuscular vaccination, followed by transfer of the APCs and soluble factors to a microfluidic human lymphoid follicle chip (LF Chip) to mimic lymphatic drainage. Non-replicating mRNA vaccines directly induce antigen expression in APCs, whereas self-amplifying mRNA vaccines require muscle cell-APC contact within the intramuscular vaccination module. Transfer of APCs and soluble factors to the LF Chip induces LF expansion, *de novo* antigen-specific IgG production against a naïve antigen (rabies virus glycoprotein), and cytokine release, with responses varying depending on LNP type. Vaccination of LF chips against SARS-COV-2 Spike recall antigen using the Moderna Spikevax vaccine generates neutralizing antibodies and induces somatic hypermutation. This biomimetic platform offers an all-human alternative for evaluating vaccine-induced immunity, potentially obviating the need for non-human primates and accelerating vaccine development.

## Introduction

mRNA vaccines have been essential in controlling the COVID-19 pandemic^1,2^. The Moderna and BioNTech mRNA vaccines against COVID-19 comprise uridine-modified, non-replicating mRNAs that encode SARS-CoV-2 spike proteins. Recently, a self-amplifying mRNA (SAM) vaccine that provides equivalent protection to non-replicating mRNA vaccines but at 6-20-fold lower doses was also approved^3–7^, further highlighting the potential of mRNA vaccine technology. However, the mechanisms underlying vaccine efficacy and toxicity have been primarily studied in inbred mice and non-human primates (NHPs), which often exhibit immune responses that differ from those seen in humans^8,9^. Furthermore, there has been an extreme shortage of NHPs for vaccine testing since the COVID-19 pandemic^10^, and testing in animals, particularly in NHPs, also raises major ethical concerns.

Although animal models have historically dominated FDA approval processes, the FDA’s new 2.0 approval pathway now expands the use of human *in vitro* data, including results generated using microfluidic human organ-on-a-chip (Organ Chip) technology, in investigational new drug (IND) applications for regulatory approval^11–13^. Despite this advance, the field of vaccine development has been slow to adopt human cell-based models. Lymphoid organoids and Organ Chip methods using human peripheral blood mononuclear cells (PBMCs) and tonsil cells have been employed for vaccine testing in academic laboratories^14–18^; however, these systems primarily focus on immune memory responses to influenza and COVID-19 vaccines and, most importantly, they have not been able to replicate clinically relevant immune responses that are generated at the injection site.

Antigen uptake, distribution, and innate immune responses are all critical factors that must be assessed to define the efficacy of mRNA vaccines^19^. However, unlike protein subunit or inactivated virus vaccines, mRNA vaccines require cellular uptake and intracellular translation of the antigen from the injected mRNA. Lipid nanoparticles (LNPs) are commonly used to encapsulate the mRNAs to promote mRNA uptake and antigen expression in human tissues, and they also may serve as adjuvants to enhance immune responses^20–23^. Consequently, there is a pressing need for human-relevant preclinical models that can more accurately study the mechanism of action of LNP-based mRNA vaccines, as well as their efficacy, toxicities, and associated biomarkers.

Most mRNA vaccines are administered through intramuscular injection. Antigen production at the muscle injection site after mRNA vaccine injection quickly peaks after 4–24 hours and then gradually wanes, and the release of pro-inflammatory cytokines such as IL-1β and IL-6 may also be important for the vaccine effects^19,24–26^. In this study, we expanded the functionality of our previously published microfluidic organ-on-a-chip model of the human lymphoid follicle (LF Chip)^16^ by combining it with a two-dimensional (2D) human intramuscular injection simulation module in which human skeletal myoblast cells (HSMCs) are cultured with antigen-presenting cells (APCs) − either monocytes or dendritic cells (DCs) − and then exposed to LNP-based mRNA vaccines *in vitro*. This module provides a method for detecting the expression of antigens encoded by mRNA vaccines and innate immune responses at the intramuscular vaccination site. By transferring cells and released factors from the intramuscular injection module to the LF Chip, adaptive immune responses can then be captured by measuring the formation of germinal center-like LFs, cytokine release, and antibody production.

Here, we used this method to compare immune responses and signaling pathways triggered by non-amplifying and self-amplifying mRNA (SAM) vaccines, as well as different LNP formulations. The method successfully modeled immune responses induced by SAM vaccines against rabies virus glycoprotein, which is a naïve antigen for most United States-based donors. Somatic hypermutation and production of cross-reactive antibodies against various SARS-CoV-2 variants with neutralization activity were also detected when the Moderna bivalent COVID-19 mRNA vaccine was administered to this *ex vivo* intramuscular vaccine model.

## Results

### Contact between APCs and HSMCs is required for modeling intramuscular vaccination

Monocytes are known to infiltrate vaccine injection sites, take up SAM vaccines, express the antigens, and transport them to the lymph nodes^19,21,27,28^. To explore modeling of intramuscular vaccination with a naïve antigen, we introduced a SAM vaccine encoding Rabies Virus Glycoprotein (RABV-G) and green fluorescent protein (GFP) encapsulated in an LNP (SAM-LNP1) into cultures of primary human monocytes or monocyte-derived DCs either when grown alone or co-cultured with HSMCs **(Fig. 1a)**. Non-toxic doses (1 to 100 ng/mL; **Supplementary Fig. 1a,b**) of SAM-LNP1 did not drive reporter expression (GFP) in monocyte monocultures (as assessed by flow cytometry), implying that the antigen was not expressed (**Fig. 1b).** However, when the SAM-LNP1 was added to co-cultures of HSMCs and monocytes **(Fig. 1b, c** and **Supplementary Fig. 1c)** or DCs **(Supplementary Fig. 2a)** to create an intramuscular simulation environment, a dose-dependent increase in GFP expression was detected in both monocytes and DCs suggesting successful antigen expression. While the GFP expression levels were variable between donors **(Fig. 1c, Supplementary Fig. 1c),** the highest level of GFP reporter expression was observed at the 100 ng/mL dose of SAM-LNP1; thus, this dose was used in all subsequent studies.

**Fig. 1.**
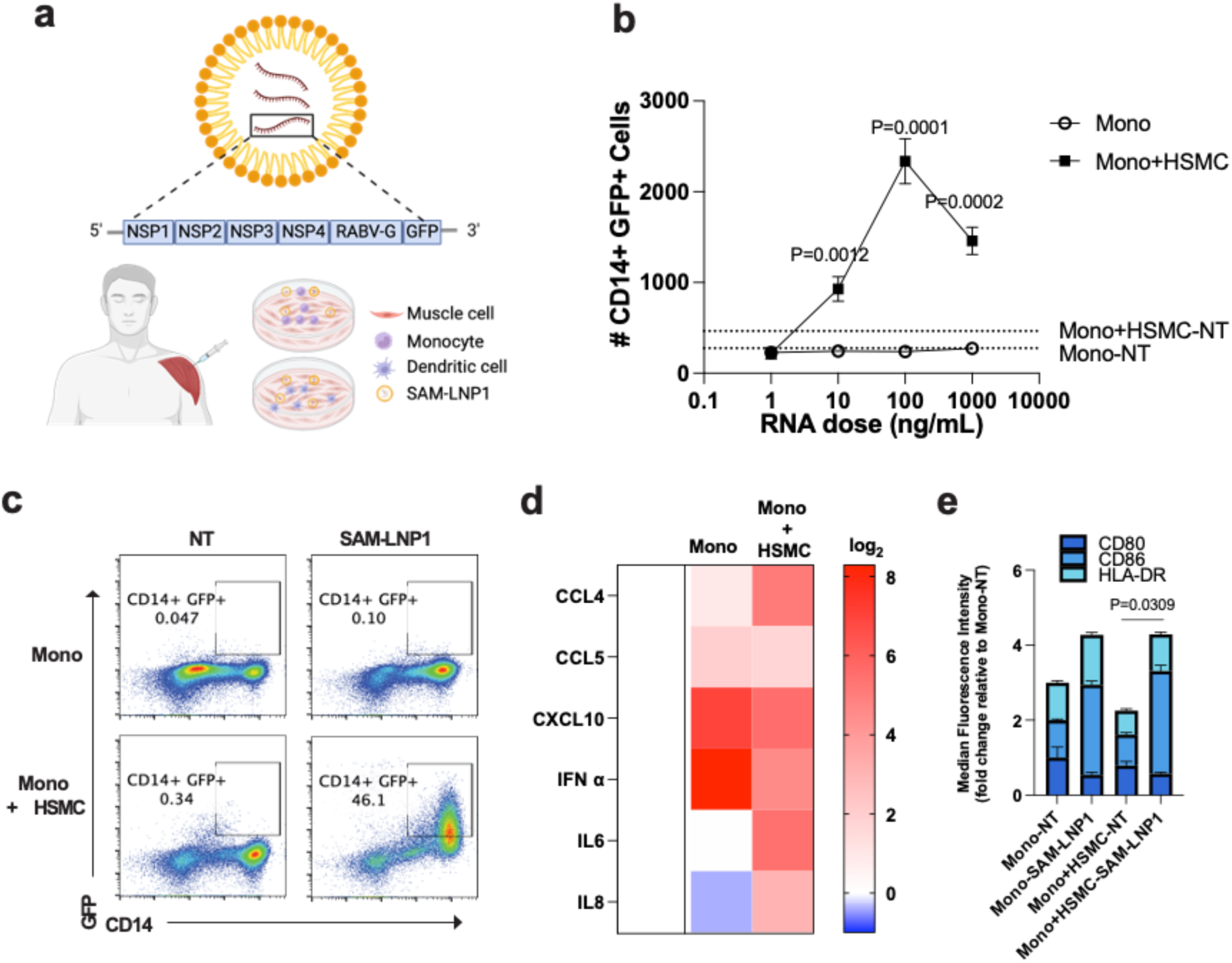
*In vitro* model of intramuscular vaccination using SAM-LNP. a, Schematic of the intramuscular vaccination model. **b,** Quantification of live CD14^+^ GFP cells by flow cytometry in monocyte (Mono) monoculture and in co-culture with HSMC (Mono+HSMC) without treatment (“no treatment”, NT) or various doses of SAM-LNP1. **c,** Density dot plots illustrating GFP expression in NT and 100ng/ml SAM-LNP1treated groups. **d,** Cytokine levels detected using a Meso Scale Discovery assay, comparing NT and 100 ng/ml SAM-LNP1in Mono and Mono+HSMC conditions. Fold change (log2) in SAM-LNP1 treated cultures relative to NT cultures is displayed. **e,** Detection of myeloid activation markers CD80, CD86, and HLA-DR by flow cytometry, comparing NT and 100 ng/ml SAM-LNP1in both Mono and Mono+HSMC settings. Representative results from one donor are shown, with similar outcomes observed in at least three donors. In **b** and **e**, mean values from 3 wells are shown. Error bars indicate standard deviation (SD). An unpaired t-test was performed in panel **b**, and two-way ANOVA in panel **e**.

Upon SAM-LNP1 vaccination, increases in levels of multiple inflammatory cytokines were observed in both monocyte monocultures and the combined cultures; however, levels of CCL4, IL-6, and IL-8 were notably higher in the co-culture model when HSMCs were present **(Fig. 1d)**. These cytokines are known to support adaptive immune responses. SAM-LNP1 treatment also increased the expression of CD86 and HLA-DR, which are markers of monocyte and DC activation **(Fig. 1e** and **Supplementary Fig. 2b).** However, CD80 was not induced; CD80 is often upregulated later or under specific conditions, such as prolonged inflammation^22,29^. These results highlight the ability of HSMCs to promote antigen expression and activation of antigen presenting cells and demonstrate the value of the co-culture model.

To further investigate the role of HSMCs in antigen expression by APCs, we conducted co-cultures in Transwell^TM^ (TW) inserts with varying pore sizes (0.4, 3, or 8 µm) placed above a HSMC monolayer **(Supplementary Fig. 3a)**. As expected, both untreated and SAM-LNP1 vaccinated monocytes were able to migrate from the upper well into the lower chamber containing the HSMCs when they were cultured on filters containing 3 or 8 µm pores, but not the 0.4 µm pore, and vaccinated cells migrated to a greater degree **(Supplementary Fig. 3b)**. Importantly, only vaccinated monocytes that were able to migrate through the larger pores and come in direct contact with the HSMCs displayed expression of GFP-labeled antigen and no antigen reporter expression was observed in monocytes that remained on top of the filter membrane in either the untreated or vaccinated cultures **(Supplementary Fig. 3c)**. These findings indicate that direct cell-cell contact between HSMCs and monocytes is necessary for antigen expression by monocytes vaccinated with SAM-LNP1.

### Mimicking vaccine drainage to lymph nodes from muscle using linked human LF Chips

To mimic the drainage of the vaccine and migration of monocytes from the vaccine site to the lymph node, we removed the conditioned medium from the co-culture of monocytes and HSMCs and flowed it through the top flow channel of a 2-channel, human LF Chip microfluidic culture device, which has been previously shown to support self-assembly of germinal center-like structures^16^. The parallel lower channel that is separated from the top channel by a porous membrane contains autologous B and T cells as well as monocytes isolated from the vaccinated intramuscular injection module embedded at a high lymph node-like density within an extracellular matrix gel **(Fig. 2a)**. When we analyzed multiple LF Chips created with cells from a single donor, we found that exposure to conditioned medium and monocytes from SAM-LNP1 vaccinated co-cultures with HSMCs resulted in a significant increase in the number of germinal center-like LFs within the chips (**Fig. 2b**). While IL-2 and IL-4 often must be added to improve T and B cell survival *in vitro*, the addition of IL-2 and IL-4 did not further enhance LF formation or increase their size in this model (**Fig. 2b,c**).

**Fig. 2.**
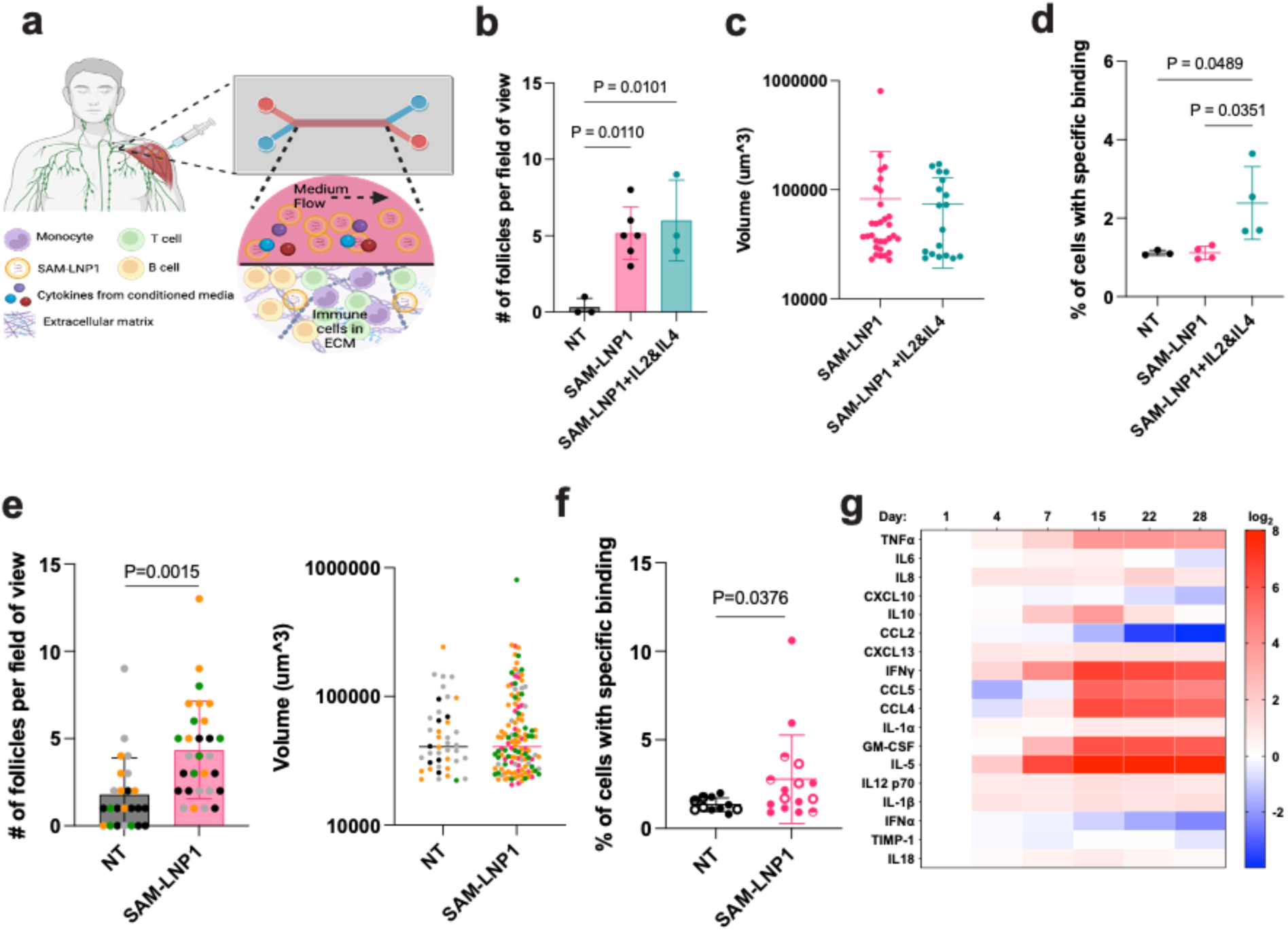
Induction of LFs and anti-RABV-G IgG production by SAM-LNP1 in human LF Chips. a, Schematic of the LF chip created with monocytes and conditioned medium from intramuscular vaccination mimicking module. Quantification of the number (**b**) and size (**c**) of LFs in LF chips of one donor based on immunostaining followed by confocal imaging. Each data point represents one field of view (**b**) or an individual follicle (**c**). **d,** Anti-RABV-G IgG levels in effluents of no treatment LF chips or chips vaccinated with or without IL2 and IL4 and cultured for 14 days were detected using a cell-based assay. **e,** Quantification of the number (left) and size (right) of LFs in LF chips from four different donors, with each color representing a different donor. Each data point corresponds to one field of view (left) or an individual follicle (right). **f,** Anti-RABV-G IgG levels in effluents of LF chips. Each data point represents one chip, with different symbols indicating chips from three independent donors. **g,** Heatmap showing Log2 fold changes in cytokine levels in LF chip effluents, measured using a Luminex® Multiplex Assay at various time points (4, 7, 15, 22, and 28 days post-vaccination) compared to day 1. Data shown are mean ± SD; **b, c, d** and **e**, One way ANOVA test, **f**, unpaired t-test.

Next, we assessed whether the SAM-LNP1 vaccinated LF Chips were able to generate antigen-specific, anti-RABV-G IgG by testing whether the chip outflows could bind to RABV-G-expressing 293T cells (**Supplementary Fig. 4**). Addition of IL-2 and IL-4 increased production of antigen-specific IgG when a single donor was analyzed (**Fig. 2d**). But when we analyzed multiple chips created with cells from 4 different human donors and treated both vaccinated and unvaccinated chips with IL-2 and IL-4, we confirmed again that addition of these cytokines induced an increase in the number of LFs, but no change in size in vaccinated chips (**Fig. 2e**). Furthermore, when we quantified anti-RABV-G IgG levels in chips created with cells from 3 donors, we found that some chips from each donor secreted anti-RABV-G IgG above baseline levels, while others did not **(Fig. 2f)**. Given the high diversity of the human immune repertoire, the variation between chips from the same donor is likely influenced by the presence and persistence of anti-RABV-G B cells. Indeed, when the B cell repertoire (BCR nucleotide sequences) from 2 different donors were compared, there was only ∼5% overlap **(Supplementary Fig. 5)**. Finally, when we compared the levels of 18 different cytokines in the LF Chip outflows during 28 days of culture, we found that cytokines known to support the germinal center response and B cell differentiation, such as IL-5, IL-6, CCL4, and IFN-γ^19,20,22,30^, increased during the first 2 weeks, followed by a subsequent plateau (**Fig. 2g**). Importantly, as these donors likely have never been vaccinated for rabies or been exposed to this antigen previously, these response are *de novo* vaccination responses.

### SAM-LNP1 and Non-Amplifying mRNA-LNP Vaccines Elicit Distinct Immune Responses

Non-amplifying mRNA vaccines, such as the Moderna and Pfizer COVID vaccines, may employ different immunological mechanisms than SAMs, and hence have different efficacies and toxicities. When we treated HSMC cocultures with monocytes or DCs with a non-amplifying, LNP-based mRNA vaccine against RABV-G antigen (NAM-LNP1), we found that, unlike the SAM vaccine, it showed higher antigen expression in DCs compared to monocytes one day after exposure in coculture with HSMC **(Fig. 3a).** Moreover, the overall expression induced by NAM-LNP1 was significantly lower than with the SAM-LNP1 (not shown) even when using a 10 times higher dose (1,000 ng/ml), likely due to its inability to amplify the response. LNPs can affect tissue tropism and reactogenicity of the vaccine cargo. Thus, we also tested another LNP (LNP2) and found that the same NAM vaccine formulated in LNP1 or LNP2 exhibited different levels of antigen presentation in DCs **(Fig. 3b)**.

**Fig. 3.**
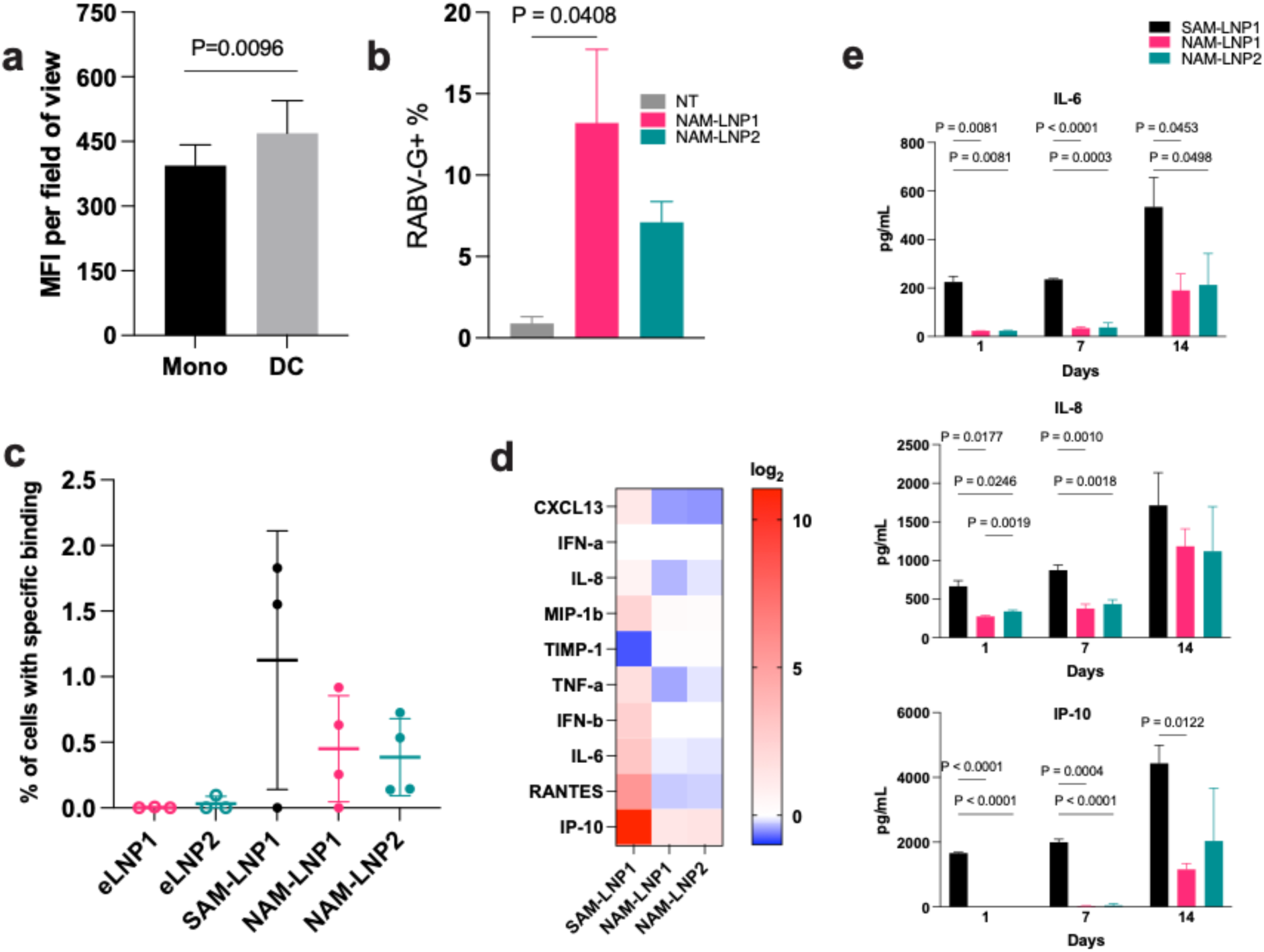
Induction of anti-RABV-G IgG and cytokine release by non-replicating mRNA. a, Quantification of monocytes and DCs after a one-day co-culture with HSMC and 1000 ng/mL of NAM-LNP1, assessed using immunofluorescence staining and detected by high-throughput imaging. Each data point represents the median fluorescence intensity (MFI) from a single field of view. **b,** Quantification of RABV-G^+^ DCs, by flow cytometry, one day after exposure to either 1000 ng/mL of NAM-LNP1 or NAM-LNP2. **c,** Anti-RABV-G IgG levels were detected in the effluents of LF chips cultured for 14 days using a cell-based assay. Each vaccine was tested in 2 donors, and similar results were obtained. **d,** Heatmap showing Log2 fold changes in cytokine levels measured via a Meso Scale Discovery assay in supernatants from HSMC-DC coculture, one day after treatment with SAM-LNP1, NAM-LNP1, and NAM-LNP2 as compared to the empty LNP control. **e,** Fold change in IL-6, IL-8, and IP-10 levels measured via a Meso Scale Discovery assay in supernatants from LF chips, 1, 7, and 14 days after treatment with SAM-LNP1, NAM-LNP1, and NAM-LNP2 as compared to the empty LNP control. Data points represent individual chips. Data shown are mean ± SD; **a,** unpaired t-test; **b** and **c**, one-way ANOVA test; **e**, two-way ANOVA test.

To test whether SAM-LNP1 vaccination (100 ng/ml) using DCs as APCs would be sufficient to induce anti-RABV-G IgG, we transferred the DC of 2 donors and the outflows from the HSMC cocultures to 3-4 LF Chips per donor and treated the chips with IL-2 and IL-4 as described above. We detected anti-RABV-G IgG in the effluents of two-thirds of the chips when SAM-LNP1 was tested using DCs as the APC. NAM-LNP1 and NAM-LNP2 (1,000 ng/ml) also induced the production of anti-RABV-G IgG compared to empty LNPs, although the levels were modest compared to those induced by SAM-LNP1 **(Fig. 3c)**. The induction of inflammatory cytokines by NAM-LNP1 and NAM-LNP2 also was significantly lower compared to SAM-LNP1, both during the first day of 2D culture and after 14 days in LF Chips **(Fig. 3d,e)**.

We then performed quantitative mass spectrometry to compare the proteomic responses induced by NAM-LNP1 (1,000 ng/ml) versus SAM-LNP1 (100 ng/ml) in immune cells extracted from vaccinated LF chips. Proteomic analysis revealed that SAM-LNP1 caused distinct and broader perturbations in the host proteome compared to NAM-LNP1 **(Fig. 4a)**. Eight proteins were downregulated by both SAM and NAM. Among these were H1.4 and H1.5 that are critical for chromatin compaction and which have been reported to influence immune cell differentiation by modulating multiple signaling pathways^31,32^. There were no common upregulated proteins between SAM and NAM-treated LF chips, suggesting significant differences in the pathways altered by SAM and NAM vaccines.

**Fig. 4.**
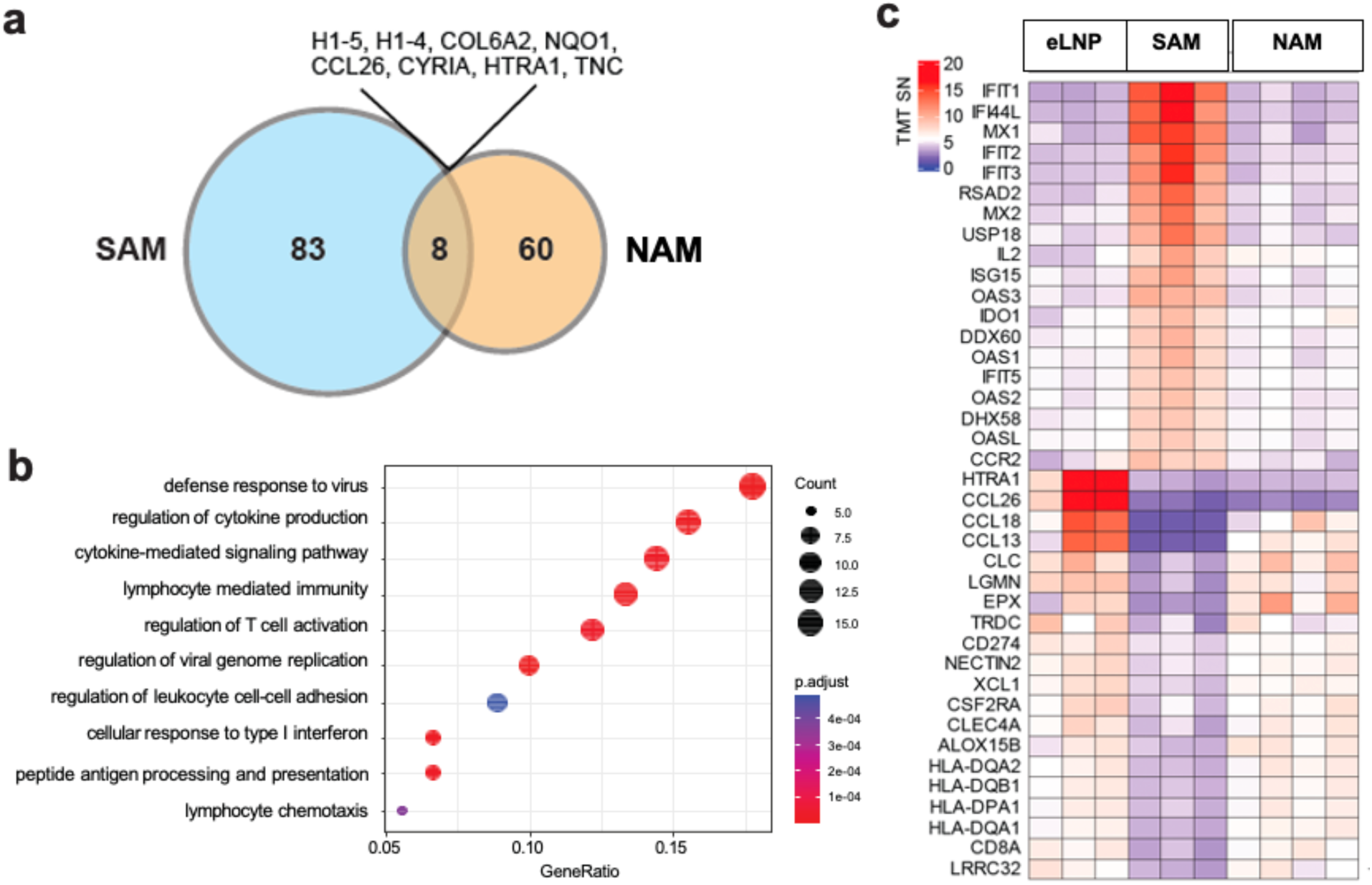
Different proteomic responses induced by self-amplifying mRNA and non-amplifying mRNA vaccines. a, Venn diagram showing the number of overlapping vs. distinct protein perturbations induced by SAM-LNP1 and NAM-LNP1. **b,** Gene ontology analysis highlights biological processes predominantly regulated by SAM-LNP1 compared to the empty LNP1. **c,** Heatmap illustrating the top differentially expressed proteins, labeled by gene locus name, showcasing the extent of regulation by the treatments.

SAM-LNP1 predominantly affected proteins involved in antiviral defense and cytokine production regulation **(Fig. 4b)**, strongly upregulating interferon response proteins, particularly members of the antiviral oligoadenylate synthase (OAS) ^33^ and interferon-induced proteins with tetratricopeptide repeats (IFIT)^34^ families. This is consistent with cellular sensing of replicating RNA. While OAS protein expression is variable, IFIT proteins are typically expressed at low levels in healthy cells and are strongly induced upon interferon exposure or viral infection. Similarly, myxovirus resistance (MX) proteins, which are antiviral proteins induced by viral infection^35^, were significantly upregulated by SAM-LNP1. Additionally, SAM vaccines induced the expression of C-C motif chemokine receptor 2 (CCR2), which plays a key role in the maturation of antigen-presenting cells^36^ and their trafficking to draining lymph nodes^37^ via the expression of C-C motif ligand 2 (CCL2, also known as monocyte chemoattractant protein 1, MCP-1). IL-2, crucial for the early priming and activation of T cells^38,39^, was also significantly induced by SAM vaccines.

In contrast, NAM-LNP1 induced only modest proteomic changes compared to the empty LNP group **(Fig. 4c, Supplementary Fig. 6)**, consistent with the significantly lower mRNA levels^40^ in the NAM versus SAM groups. Notably, the proteins upregulated by NAM-LNP1 are closely associated with T cell responses. Among them, Kinase Suppressor of Ras 1 (KSR1), a scaffold protein, plays a crucial role in regulating the activation threshold of Extracellular Signal-Regulated Kinase (ERK) in T cells^41,42^; Unc-13 Homolog D (UNC13D) is essential for mediating the release of cytolytic granules from T cells^43^; and Fatty Acid Binding Protein 4 (FABP4) is involved in modulating T cell receptor signaling pathways^44^.

### Somatic hypermutation induced by mRNA-LNP vaccines

To study the recall responses of mRNA vaccines in this LF Chip-based intramuscular vaccination platform, we used the Moderna Spikevax bivalent Original and Omicron BA.4/BA.5 vaccine and conducted experiments using both DCs and monocytes from vaccinated HSMC cocultures as well as conditioned medium along with IL-2 and IL-4, using the linking method described above. This vaccine was chosen because its induction of neutralizing antibodies has been well-documented in humans, and somatic hypermutations (SHMs) often require multiple vaccine exposures ^45^. In multiple LF chips created from the cells of 4 different donors, vaccination using the Moderna mRNA vaccine (500 ng/ml) induced robust IgG responses against the SARS-CoV-2 Wuhan strain, as well as the Omicron BA.4 and BA.5 variants. The IgG titer in unvaccinated and vaccinated chips against the Wuhan spike was 3 to 10 times higher than the titer against the BA.4 and BA.5 spikes **(Fig. 5a),** which may reflect the wide distribution of the monovalent vaccines against the Wuhan strain within Massachusetts where our immune cell donor pool is located. We also verified the neutralization efficiency of the antibodies produced by the LF Chips from 2 donors **(Fig. 5b)** by testing the ability of the chip outflows to block infection of ACE2-expressing 293T cells by a pseudovirus expressing the Wuhan strain spike protein.

**Fig. 5.**
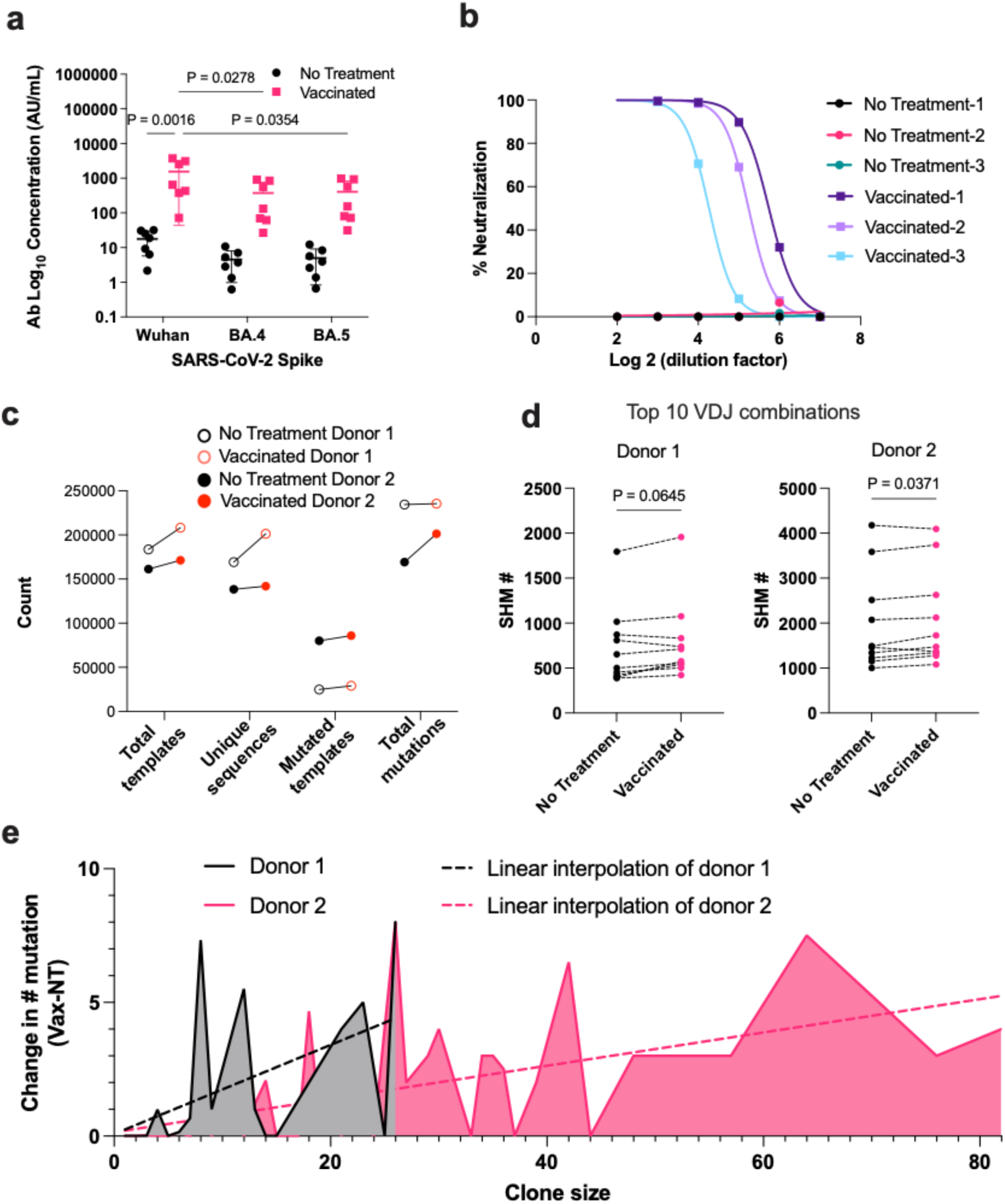
Induction of spike-specific neutralizing antibodies and somatic hypermutation in LF chips following vaccination with Moderna bivalent COVID-19 vaccine. a, Moderna bivalent COVID-19 vaccine was administered at 500 ng/ml (M500). Day 9 LF chip outflows post-vaccination (M500) and no treatment (NT) were tested to determine COVID-19 spike-specific IgG concentrations using the Meso Scale Discovery (MSD) V-PLEX SARS-CoV-2 Panel 27 (IgG) Kit. **b,** Neutralization by outflows from each no treatment or vaccinated chip created from the cells of one individual donor is depicted. 0% neutralization is defined by the luciferase signal in cells infected with SARS-CoV-2 pseudovirus using fresh medium controls, and 100% neutralization corresponds to the signal in uninfected cells. Similar results were obtained from a 2^nd^ donor’s cells. **c,** BCR repertoire analysis on LF chips from two donors. Total templates, unique sequences, mutated templates, and total mutations of DNA templates from cells recovered from both vaccinated (M500) and no treatment chips from two donors are shown. **d,** Comparison of SHM counts of top 10 V-D-J combinations from both vaccinated (M500) and no treatment chips from both donors. **e,** Changes in average SHM in vaccinated samples compared to unvaccinated samples across both donors. Data shown are mean ± SD, **a,** two-way ANOVA test, **d,** Wilcoxon matched pairs signed rank test.

To understand whether the variable neutralization efficiency of the induced antibodies in the outflow from different chips and donors **(Fig. 5b)** reflects a lack of B cell activation and maturation or the detection limits of our workflow, we sequenced the CDR3 loop of the IgH locus from cells isolated from the LF Chips from 2 donors 14 days after vaccination. Due to the diversity of the human immune repertoire and the limited overlap between chips from the same donor, sequences from 3 chips for each donor were pooled for further analysis; over 100,000 unique sequences were obtained per donor (**Fig. 5c**). Both donors showed an increase in the total number of B cell sequences, unique sequences, B cells with SHMs, and total SHMs upon vaccination. However, donor 2, which produced higher levels of neutralizing antibodies, had more B cells with existing mutations, and yet a significant increase in SHM was still observed upon vaccination; vaccination did not alter V, D, or J gene usage (data not shown).

The top 10 V-D-J combinations had >100 mutations in the LF Chip even without vaccination, which indicates the baseline diversity of the donor BCR repertoire, but vaccination resulted in 5-10% increase in SHMs in donor 2 (**Fig. 5d**), and a similar trend was observed in donor 1. We found that vaccinated chips had a higher average level of SHMs as clone size increased **(Fig. 5e)**, a pattern evident in both donors, and donor 2 had larger clones than donor 1. To look at these effects in greater detail, we compared sequences from specific, highly expanded V-D-J combinations between vaccinated and unvaccinated chips. As shown for a representative V-D-J combination from donor 2 (**Fig. 6)**, vaccination increased the number of mutated sequences and the number of mutations in expanded clones, confirming that the LF chip displays SHM upon stimulation with vaccines.

**Fig. 6.**
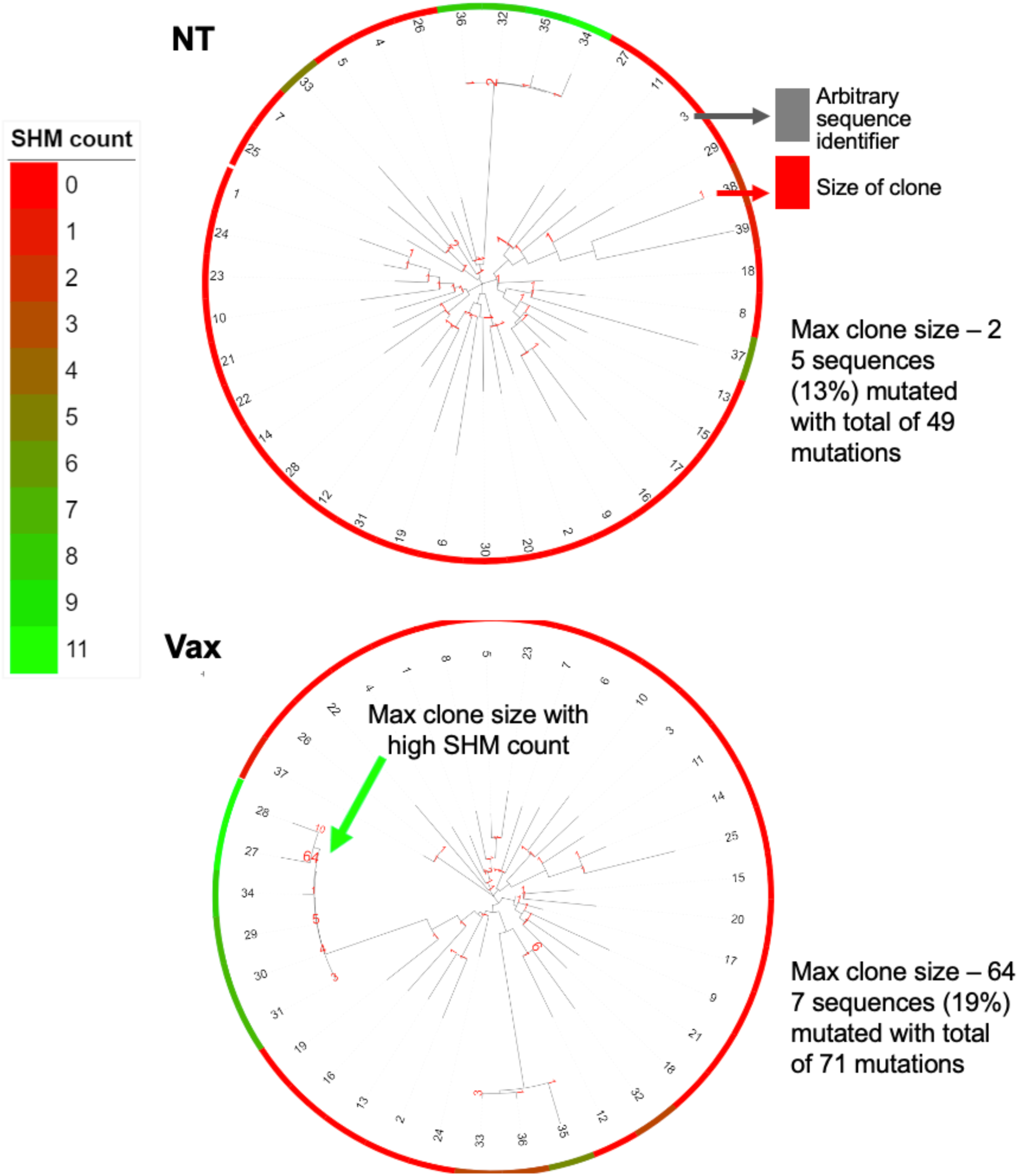
Clonal Expansion and Somatic Hypermutation in LF Chips. Clonal sequences of BCR containing the V-D-J genes IGHV01-69, IGHD04-23*01, and IGHJ06- 01*02 were analyzed using ClustalW for alignment. Phylogenetic analysis was subsequently visualized with iTol, generating the depicted dendrograms. **a,** Dendrogram of sequences from no treatment chips. **b,** Dendrogram of sequences from chips vaccinated with 500ng/ml Moderna bivalent COVID-19 vaccine. Colors on the outer ring indicate the number of mutations per sequence. Branch labels represent the size of each clone, and the numbers within the ring serve as arbitrary sequence identifiers.

## Discussion

The efficacy of mRNA vaccines is closely tied to the mRNA format, delivery vehicle, and route of administration^19,46,47^. In this study, we developed an *ex vivo* model mimicking the intramuscular injection site and combined it with LF Chips to create an experimental platform that provides a comprehensive view of mRNA vaccine uptake and immune responses in humans. We observed that the expression of antigen encoded by SAM vaccine in the APCs depends on co-culture with muscle cells and the formation of direct cell-cell contacts. This co-culture also enhanced the activation of APCs in response to mRNA vaccination, as indicated by increased expression of surface markers (e.g., CD86, HLA-DR), and the release of pro-inflammatory cytokines, including IL-6 and IL-8. Interestingly, IL-6, a key regulator of T follicular helper cells^48–50^, was specially induced by SAM vaccines when APCs were co-cultured with muscle cells. These findings emphasize the importance of studying vaccine distribution, innate immune responses, and the contribution of stromal cells to responses, particularly to SAM vaccines, as previously noted by others^51,19,52,45,53^. In contrast, NAM vaccines can be taken up directly by monocytes and DCs in the absence of muscle cells; however, the LNP component affects cytokine release and antigen expression^17,21,51,54^.

To detect adaptive immune responses induced by mRNA vaccines, we mixed APCs from the intramuscular vaccination simulation module with autologous T and B cells to create an LF Chip model of a draining lymph node, leveraging our published method^16^. The conditioned medium from the intramuscular vaccination module containing released soluble factors was also transferred to the top channel of the LF Chip. Notably, we found that SAM vaccines induce LF formation, and the addition of IL-2 and IL-4 enhances antigen-specific antibody production, although it does not further alter the number or size of the LFs. These LF Chips can be maintained for up to 28 days in culture, allowing for the long-term study of cytokine release induced by mRNA vaccines. The secreted cytokines we detected, including IL-1α, IL-1β, IL-6, and CXCL10, are consistent with findings from previous *in vivo* studies of SAM vaccines^20,22,23,30^. Our experiments also confirm that the LF Chip supports immunoglobulin G generation when challenged with naïve antigens, a capability lacking in other systems that utilize blood-derived cells^17,18^. Notably, only approximately 0.0055% to 0.088% of B cells in peripheral blood bind RABV-G^55^. Given that Rabies patients were excluded from our donor pool and that RABV-G shares only ∼20% sequence identity even with other rhabdoviruses^56^, the responses observed in the chips likely originate from naïve B cells.

Comparing multiple LNP delivery vehicles and RNA payload types opens opportunities for *in vitro* comparative studies to identify optimal combinations of delivery vehicles and mRNA formats. Our on-chip study confirmed previous reports that SAM vaccines can induce similar immune responses at doses 10 times lower than those required for NAM vaccines^57^. Interestingly, while the cytokine induction patterns of the three mRNA vaccines tested were similar, SAM vaccines triggered significantly more robust cytokine responses and antigen-specific antibodies than NAM vaccines. This may be related to our finding from our proteomic analysis, which revealed that SAM and NAM vaccines activate distinct signaling pathways. Of the proteins altered differently by SAM and NAM vaccines, only 8 overlapped between the 2 vaccine formats, with 83 unique proteins induced by the SAM and 60 by NAM vaccines. SAM vaccines appeared to activate more robust viral defense and cytokine regulation pathways. To our knowledge, this is the first reported proteomic comparison of human immune cell proteome changes induced by different mRNA vaccine formats, and we anticipate that further studies of vaccine-induced signaling pathways will provide valuable insights for future vaccine design.

We also tested the Moderna bivalent COVID-19 mRNA vaccine to investigate memory responses and booster effects using our method for modeling intramuscular vaccination. Since most of our immune cell donors had likely received a COVID-19 mRNA vaccine due to state policies in Massachusetts, we anticipated observing memory immune responses against the SARS-CoV-2 antigen contained with this vaccine in cells from these donors. Our results were consistent with previous findings, showing that the bivalent COVID-19 mRNA vaccine induced 3- 10 times more antibodies against the Wuhan strain compared to the Omicron BA.4 and BA.5 variants^58,59^. Also, the neutralizing efficacy of the antibodies produced by the LF Chip platform was confirmed, and SHM analysis revealed that B cells underwent SHM within the chip environment. This is also the first report of inducing somatic hypermutation in blood-derived B cells *in vitro* with a clinical mRNA vaccine.

In conclusion, this study establishes a robust method for interrogating human immune responses to mRNA-LNP vaccination at both the vaccine injection site and within lymphoid tissues. In contrast to previous methods that primarily focused on memory responses^15–18^, this preclinical model is capable of evaluating both *de novo* and memory human B cell responses to multiple mRNA vaccine formats. We also characterized the dynamics of cytokine release and antibody production associated with different mRNA vaccine modalities, which can be further leveraged to profile *in vivo* immune responses during clinical trials. While this study primarily focused on cytokine release and B cell responses, future work could expand on these findings by investigating antigen-specific T cell responses. Additionally, this system could be used to study the roles of other immune cells during vaccination, such as mast cells, plasmacytoid dendritic cells, and natural killer cells. Further refinements could include adding stromal cells to the intramuscular module to accurately represent the complexity of the injection site, as fibroblasts and other stromal cell types play a crucial role in antigen uptake and innate immune responses.^60,61^ Most importantly, these data demonstrate that this experimental model of human vaccination offers a new and exciting alternative to NHP studies in future vaccine development programs.

## STAR★Methods

### Expansion of HSMCs

HSMCs (Lonza, CC-2580) were rapidly thawed in a water bath and resuspended in Skeletal Muscle Growth Medium-2 (SkGM-2, Lonza, CC-3245). Cells were seeded in a T-150 flask for expansion. Confluency and morphology were monitored daily using brightfield microscopy, and media were changed every three days. Cells were used for vaccination experiments upon reaching 70-80% confluency.

### Peripheral Blood Mononuclear Cells (PBMC) Isolation and DC Differentiation

De-identified human patient-derived apheresis collars were obtained from the Crimson Biomaterials Collection Core Facility, under Institutional Review Board approval from Harvard University (protocol 22470). PBMCs were isolated by density centrifugation using Lymphoprep (StemCell Technologies, 07801), followed by magnetic bead-based negative selection for B cells (StemCell Technologies, 19054), T cells (17951), and monocytes (19058). For cryopreservation, cells were frozen in Recovery Cell Culture Freezing Medium (Gibco, 12648-010) at 20 million cells/mL. Monocytes were differentiated into DCs by culturing in complete RPMI with 400 ng/mL GM-CSF (Mitenyi Biotec, 130-095372) and 250 ng/mL IL-4 for 5-6 days, with 50% medium replacement every 2 days.

### Rabies Virus Glycoprotein and Enhanced Green Fluorescent protein (eGFP) mRNA Preparation

The antigen sequence of the full-length Rabies Virus Glycoprotein is based on the Pasteur strain (GenBank accession number: AAA47218.1). The Open Reading Frame (ORF) was optimized for expression in mammalian cells and cloned into SAM or mRNA backbones using standard molecular biology techniques. In the SAM construct, the Rabies G gene was cloned in the first position and the eGFP (GenBank AAB02572) second, separated by the IRES from Enterovirus 71 (EV71). In vitro RNA transcription, purification, and LNP formulation were performed as described previously^62–64^. Tri-link prepared conventional mRNA with the same Rabies Virus Glycoprotein sequence, incorporating n-1-methyl-pseudouridine in place of uridine.

### LNP Formulation Preparation

Lipid and RNA components were prepared in ethanol and citrate buffer, respectively. LNPs encapsulating RNA were formulated by lipid nanoprecipitation using a modified ethanol dilution process. After the formation of the LNP, the LNP was buffer exchanged using a desalting column into a holding buffer. A sucrose excipient was added to the LNPs for final storage.

### Co-culture of Muscle Cells and Antigen-Presenting Cells (APCs) with Vaccination

Skeletal muscle cells were trypsinized using 0.05% trypsin-EDTA (Gibco, 25300054), resuspended in SkGM-2 medium, and seeded at a density of 120,000 cells per well in a 24-well plate. Cells were incubated overnight to allow for adherence. Monocytes were thawed, or cultured dendritic cells (DCs) were collected and resuspended at a concentration of 0.5 million cells/mL, and 0.5 million monocytes or DCs were added to the muscle cell wells along with the indicated mRNA vaccine.

### Human LF Chip

Two-channel microfluidic Organ Chip devices and ZOE-ORB instruments were obtained from Emulate Inc. The Organ Chips were plasma-activated and seeded as previously described^16^. Briefly, chips were perfused with 1% 3-Aminopropyltrimethoxysilane (Sigma, 281778) in ethanol for 1 hour, followed by washing the channels with ethanol. Chips were incubated in an 80°C oven overnight. For muscle-APC co-culture experiments, media were collected and pooled, and cell counts were performed. For RABV-G mRNA vaccine experiments, APCs were mixed with T and B cells at a 5: 47.5: 47.5 ratio to achieve a final concentration of 1.5 × 10^8 cells/mL in complete RPMI supplemented with Matrigel (60%; Corning, 356234) and type I collagen (15%; Corning, 354249). For COVID-19 mRNA vaccine experiments, DCs and monocytes from muscle co-cultures were mixed with T and B cells in a 1:1:49:49 ratio. A 15 µL aliquot of the cellularized ECM mixture was introduced into the bottom channel of the 2-channel Organ Chip. After allowing the matrix to gel for 30 minutes at 37°C, the top channel was filled with conditioned medium, prepared as a 1:2 ratio of co-culture to fresh media for RABV-G mRNA vaccines, and a 1:1:8 ratio of DC co-culture medium, monocyte co-culture medium, and fresh media for COVID-19 mRNA vaccines. The chips were perfused with indicated media at 40 µL/h using ZOE instruments in a 37°C incubator.

### Immunostaining and Imaging

Chips were fixed by adding 200 µL of 4% paraformaldehyde to one side and allowing 30-50 µL to flow out. Both ports were then plugged, and an additional 200 µL was added. After 1 hour of rocking at room temperature to facilitate back and forth flow of paraformaldehyde solution in the channel, chips were washed twice with PBS for 15 minutes. Nuclear staining was performed using Hoechst (1:1000 dilution, Invitrogen, H3570) for 1 hour, followed by three 15-minute PBS washes. Imaging was performed on a Leica SP5 confocal microscope, and follicle size and number were quantified using Imaris software (Bitplane, version 9.3).

For imaging vaccinated HSMC in **Supplementary Figure 1c**, HSMC vaccinated for 24 hours in a 96-well plate were imaged with a Sapphire Biomolecular Imager (Azure Biosystems) and analyzed using Sapphire software (V1.12.0921.0). In Figures 4a, co-cultured vaccinated APCs were immunostained with anti-Rabies-G (Millipore Corp, MAB8727, 1:1000) and goat anti-mouse IgG (H&L) Cross-Adsorbed Secondary Antibody (Invitrogen, A-21235, 1:2000). Images were taken with an ImageXpress Micro4 (Molecular Devices) and analyzed using MetaXpress (6.5.3.427) and ImageJ software (v 2.14.0)

### Extraction of Live Cells from Organ Chips

The gel from the bottom channel of the LF chip was collected in Cell Recovery Solution (Corning, 354253). The collection tube was centrifuged at 300g for 5 minutes, and the supernatant was aspirated. The pellet was resuspended in 1 mL of fresh Cell Recovery Solution and incubated on ice for 1 hour. Following incubation, the sample was centrifuged again at 300g for 5 minutes, the supernatant was aspirated, and the cells were prepared for downstream applications.

### Flow cytometry

Cell viability was assessed using ViaKrome 808 Fixable Viability Dye (Beckman Coulter, C36628, 1:100 dilution). Cells were stained with antibodies for CD14 (BD 563561), CD1c (BioLegend 331526), HLA-DR (BioLegend 361610), CD80 (BioLegend 305231), CD86 (BioLegend 305430), anti-Rabies-G (Millipore Corp, MAB8727) and goat anti-mouse IgG (H&L) Cross-Adsorbed Secondary Antibody (Invitrogen, A-21235). Data were collected on a CytoFlex LX (Beckman Coulter) and analyzed using FlowJo V10 software (FlowJo LLC).

### Cell-Based Assay for Anti-Rabies Antibody Detection

293T cells (1 x 10^6) were seeded in 6-well plates in DMEM with 10% FBS and 1% P/S, 48 hours prior to transfection. Cells were transfected with 50 ng/mL of SAM-RABV-G-GFP-LNPs or empty LNPs (control) and incubated in 2 mL of serum-free DMEM. We used the SAM vaccine to make RABV-G expressing 293T. Cells were incubated with outflows to be tested for anti-RABV-G antibodies and then cells stained with ViaKrome 808 dye and anti-human IgG APC (Biolegend 410712). Differential staining with anti-human IgG of control vs. RABV-G expressing 293T indicated the presence of anti-RABV-G antibodies. Specific binding was calculated as shown in **Supplementary Fig. 4**.

### Proteome Analysis by Mass Spectrometry

The mass spectrometry-based multiplexed proteomic analysis was performed as previously described^65^. Briefly, human immune cells were cultured for two weeks at LF chips with different mRNA vaccines. The cells were then lysed in 8 M urea and 200 mM EPPS pH 8.5 with Pierce Protease and Phosphatase Inhibitor (Thermo Fisher, A32961). Protein concentration was measured by BCA assay. ∼20 µg proteome in 20 µL was treated sequentially with 20 mM TCEP and 200 mM IAA for 15 minutes and 30 minutes, respectively. 3 µL SP3 beads (1:1 mixture of hydrophobic and hydrophilic type, 50 mg/mL, (Cat. #45152105050250 and Cat. #65152105050250) and 30 µL ethanol supplemented with 20 mM DTT were added to the lysate. SP3-bind proteome was washed twice with 80% Ethanol followed by sequential digestion with 0.2 µg Lys-C and 0.2 µg trypsin for 2 hours and overnight at 37 °C, respectively. The digested peptides were added 10 µL acetonitrile and labeled with 4 µL TMTpro reagent for 1 hour before combining all samples into one single multiplexed sample and desalting through Sep-Pak C18 column. The eluent was dried and then fractionated by basic-pH HPLC for total 24 fractions.

Both fractionated proteome samples were analyzed by liquid chromatography-tandem mass spectrometry (LC-MS/MS) on an Orbitrap Eclipse mass spectrometer coupled to a Proxeon NanoLC-1200 UHPLC50 and FAIMS Pro device. Data were collected alternating between a set of three FAIMS compensation voltages (CVs) at-40,-60, and-80 V was used. MS1 scans were collected in the Orbitrap with a resolution setting of 60 K for proteome, a mass range of 400-1600 m/z, an AGC at 100%, and a maximum injection time of 50 ms. MS2 scans were acquired in Top Speed mode with a cycle time of 1 s. Peptide precursors were selected and fragmented using HCD with a collision energy of 36. MS2 scans were collected in the Orbitrap with a resolution of 50K, a fixed scan range of 110-2000 m/z, and a 500% AGC with a maximum injection time of 86 ms. Dynamic exclusion was set to 90 s with a mass tolerance of ± 10 p.p.m.

The mass spectra were processed using a Comet-based pipeline and searches were performed using a 50-ppm precursor ion tolerance for the analysis of the total protein levels^66^. The product ion tolerance was set to 0.9 Da. Tandem mass tags on lysine residues and peptide amino termini (+304.2071 Da) and carbamidomethylation of cysteine residues (+57.021 Da), were set as static modifications, whereas oxidation of methionine residues (+15.995 Da) was set as a variable modification. Peptide-spectrum matches were adjusted to a 1% false-discovery rate^67^. Proteins were quantified by summing the reporter-ion counts across all matching peptide-spectrum matches, as described previously. Gene Ontology analysis and graphs was generated using R packages ClusterProfiler^68^, Complexed Heatmap and Ehanced Volcano.

### Proteomic Data Availability

The mass spectrometry data have been deposited at the ProteomeXchange Consortium and are publicly available as of the date of publication. The accession number is PXD056699. (The reviewer access can be provided if required during submission.)

### Immunoglobulin and Cytokine Quantification

To measure immunoglobulins targeting multiple SARS-CoV-2 spike protein variants in the day 9 outflow from both unvaccinated and vaccinated LF chips, the Meso Scale Discovery (MSD) V-PLEX SARS-CoV-2 Spike Panel 27 (IgG) Kit (K15437U) was used, with data acquired on an MSD Meso Sector S 600 microplate reader and analyzed with Discovery Workbench 4.0 software. The MSD V-PLEX SARS-CoV-2 Spike Panel 27 (IgG) Kit included 96-well plates immobilized with Wuhan, BA.4, and BA.5 spike antigens (data reported in Fig. 6a), along with a beta variant (B.1.351), delta variant (B.1.617.2; AY.4 Alt Seq 2) and additional omicron sub-lineage spike proteins (BA.2, BA.2.12.1, BA.2+L452M, BA.2+L452R, and BA.3; data not shown), in addition to a reference standard and three controls with known levels of antibodies against antigens in the kit. Assays were performed following the MSD-provided, kit-specific protocol, testing LF chip outflows in duplicate and including CH65 IgG monoclonal antibody (mAb) (influenza receptor binding site-specific mAb) as a negative control. Data from LF chip outflow dilutions (1:10, 1:70, 1:490) falling nearest to the midpoint of the accurate range of the standard curve were selected for reporting. Values chosen were back calculated against the standard curve and reported as arbitrary units per milliliter (AU/mL), accounting for dilution factor.

The cytokine levels in the media were assessed using a custom MSD kit (as presented in **Figures 1d** and **3d,e**) and read using an MSD microplate reader model 1300 alongside the Discovery Workbench 4.0 software. Customized Luminex® Multiplex Assay kits from Thermo Fisher Scientific were employed for the cytokine concentrations in the effluents depicted in **Figure 2g**. Analyte concentrations were determined on a Bio-Plex 3D suspension array system, and the standard curve fitting and concentration calculations were conducted using Bio-Plex Manager software (Bio-Rad, version 6.0).

### BCR Nucleotide Sequences Analysis

Human immune cells were cultured on LF chips for two weeks with 500 ng/ml of the Moderna bivalent COVID-19 mRNA vaccine. Following this, cells were extracted from the chips and sent to Adaptive Biotechnologies for sequencing of the CDR3 loop of IgH. The data was then analyzed and visualized using the immunoSEQ Analyzer, Clustal W^69^, iTOL^70^ and pivot charts in Prism 10.

### Statistics

All experiments (excluding proteomics) were performed at least twice, with at least three technical replicates for each group in each experiment. Data are presented as mean values ± standard deviation (SD). Graphing and statistical comparisons were carried out using GraphPad Prism software (v10.1.0). Statistical significance for comparisons between two groups was determined using an unpaired t-test. A P-value of <0.05 was considered statistically significant, with exact P-values labeled in all figures.

## Acknowledgments

The studies on SAM and NAM vaccines were sponsored by GSK. The studies with Moderna vaccines and on somatic hypermutation were supported by funding from the Bill and Melinda Gates Foundation (INV-002274 to G.G. and D.E.I) and the in-kind Women in Science Award from Adaptive Biotechnologies (to G.G.), Development of the LF chip was also supported by funding from the Bill and Melinda Gates Foundation (INV-002164 to D.E.I), the Biomedical Advanced Research and Development Authority (contract 75A50121C00075 to D.E.I) and the Biomedical Advanced Research and Development Authority-U.S. Food and Drug Administration (contract 75A50123D00004 to D.E.I), SARS-CoV-2-specific IgG antibody quantification in organ-chip outflows supported by funding from Gates Foundation (INV-060822 to G.D.T.), as well as by a Wyss Technology Development Fellowship (to Y.Z.). The authors would like to thank Harvard University Health Services for sharing the unused portion of the Moderna vaccine from opened vials to enable the studies on the production of neutralizing antibodies and somatic hypermutation.

## Author contributions

Y.Z., J.M., D.E.I, and G.G. conceived the study. J.M., D.E.I, and G.G. oversaw the study. Y.Z., M.W.K., K.Y., L.H., S.F., A.S.P., G.M., S.B., K.L., C.A.H., L.D.W., G.D.T., P.Y., S.G., J.M., D.E.I., and G.G. participated in experiment design and/or discussions. Y.Z., M.W.K., K.Y., A.P., P.P., L.H., Y.M., S.S., S.G., A.R.I., J.F., K.M., E.M., C.A.H., and L.D.W. performed the experiments and data collection. Y.Z., M.W.K., K.Y., D.E.I., and G.G. analyzed the data, prepared the figures and tables, and wrote the manuscript with input from all authors. D.E.I. and G.G. reviewed and edited the manuscript. All authors approved the final version.

## SUPPLEMENTARY FIGURES

**Supplementary Fig. 1.**
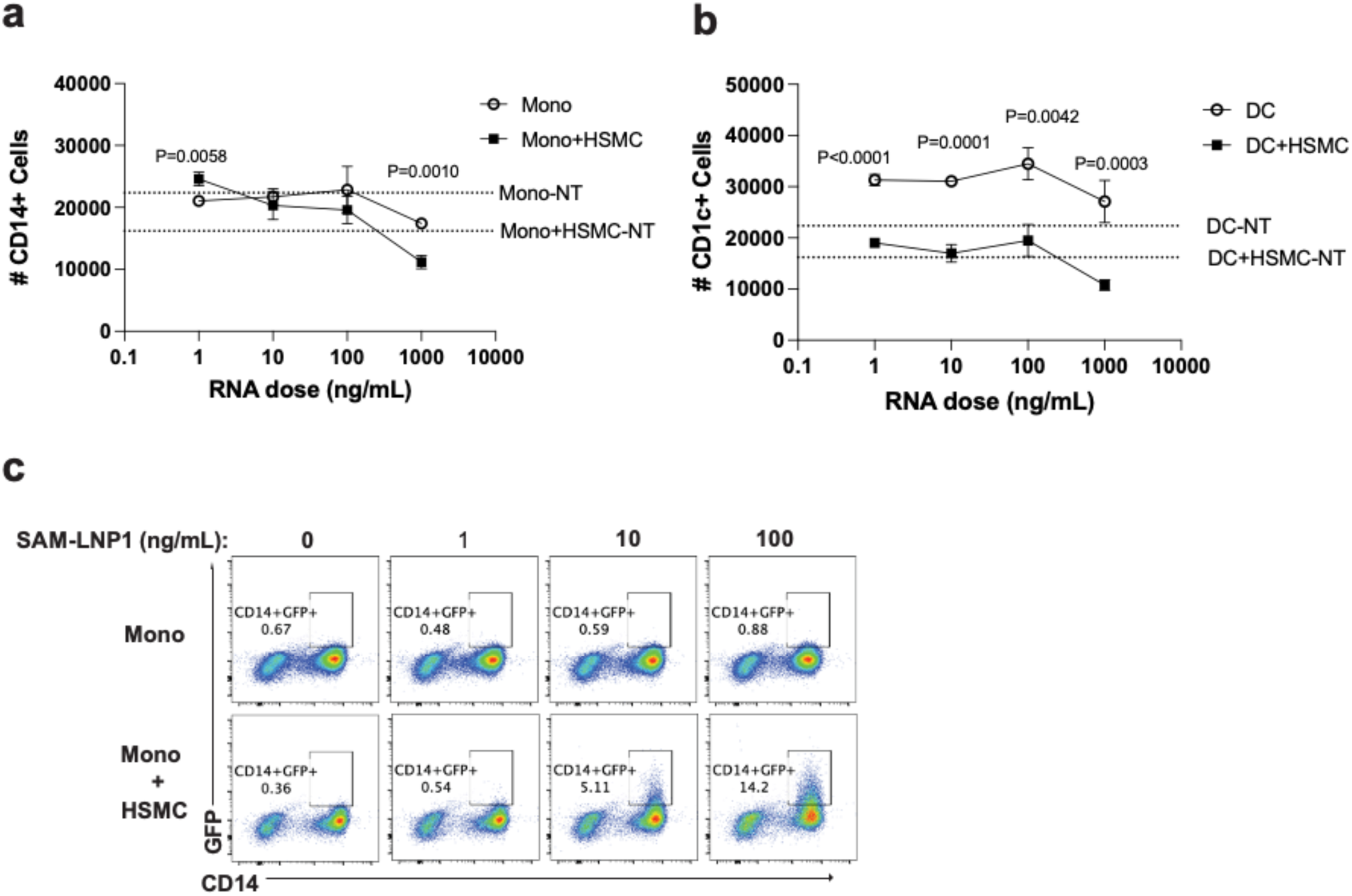
Dose-dependent effects of SAM-LNP1 on antigen-presenting cells (APCs) and human skeletal muscle cells (HSMCs). Quantification of **a,** live CD14^+^ monocytes. **b,** live CD1c^+^ DCs in monoculture and co-culture under no treatment (NT) or various doses of SAM-LNP, measured by flow cytometry. **c,** Monocyte uptake of SAM-LNP1 in co-culture with HSMCs, showing a dose-dependent pattern represented through percentage changes in flow cytometry data. Representative results from one donor, with similar results obtained in at least three donors. Data shown are mean values from 3 wells ± SD; an unpaired t-test was performed for **a** and **b.**

**Supplementary Fig. 2.**
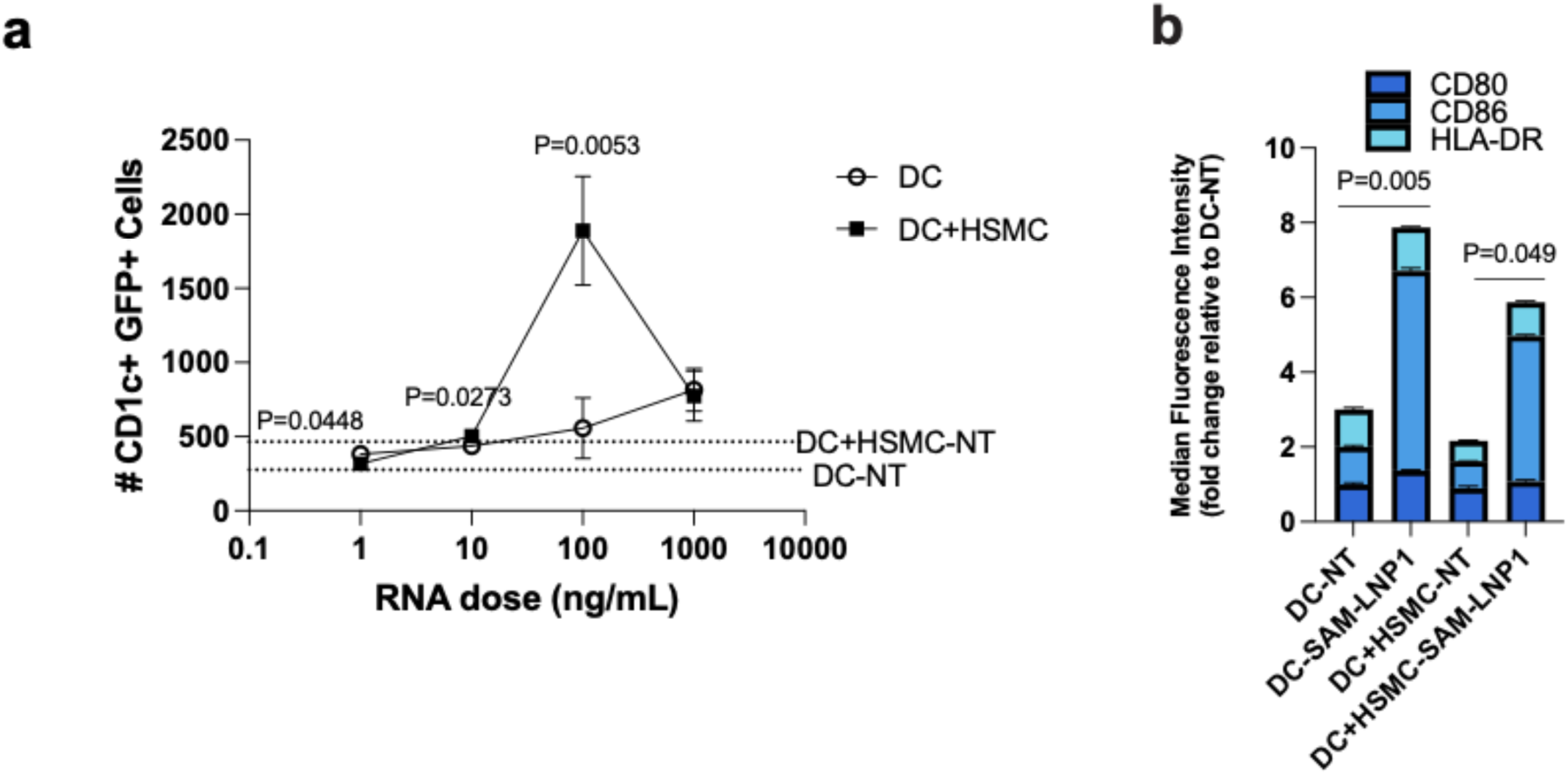
Dose-dependent effects of SAM-LNP1 on DCs and HSMCs. a, Quantification of live CD1c^+^ GFP^+^ DCs recovered from monocultures (DC) or co-cultures with human skeletal muscle cells (DC+HSMC) under no treatment (NT) or various doses of SAM-LNP1, using flow cytometry. **b,** Detection of DC activation markers CD80, CD86, and HLA-DR by flow cytometry, comparing NT to 100 ng/ml SAM-LNP1 in DC alone (DC) and DC-muscle cell co-culture (DC+HSMC) conditions. Representative results from one donor, with similar results obtained in at least three donors. Data shown are mean values from 3 wells ± SD; an unpaired t-test was performed for **a**.

**Supplementary Fig. 3.**
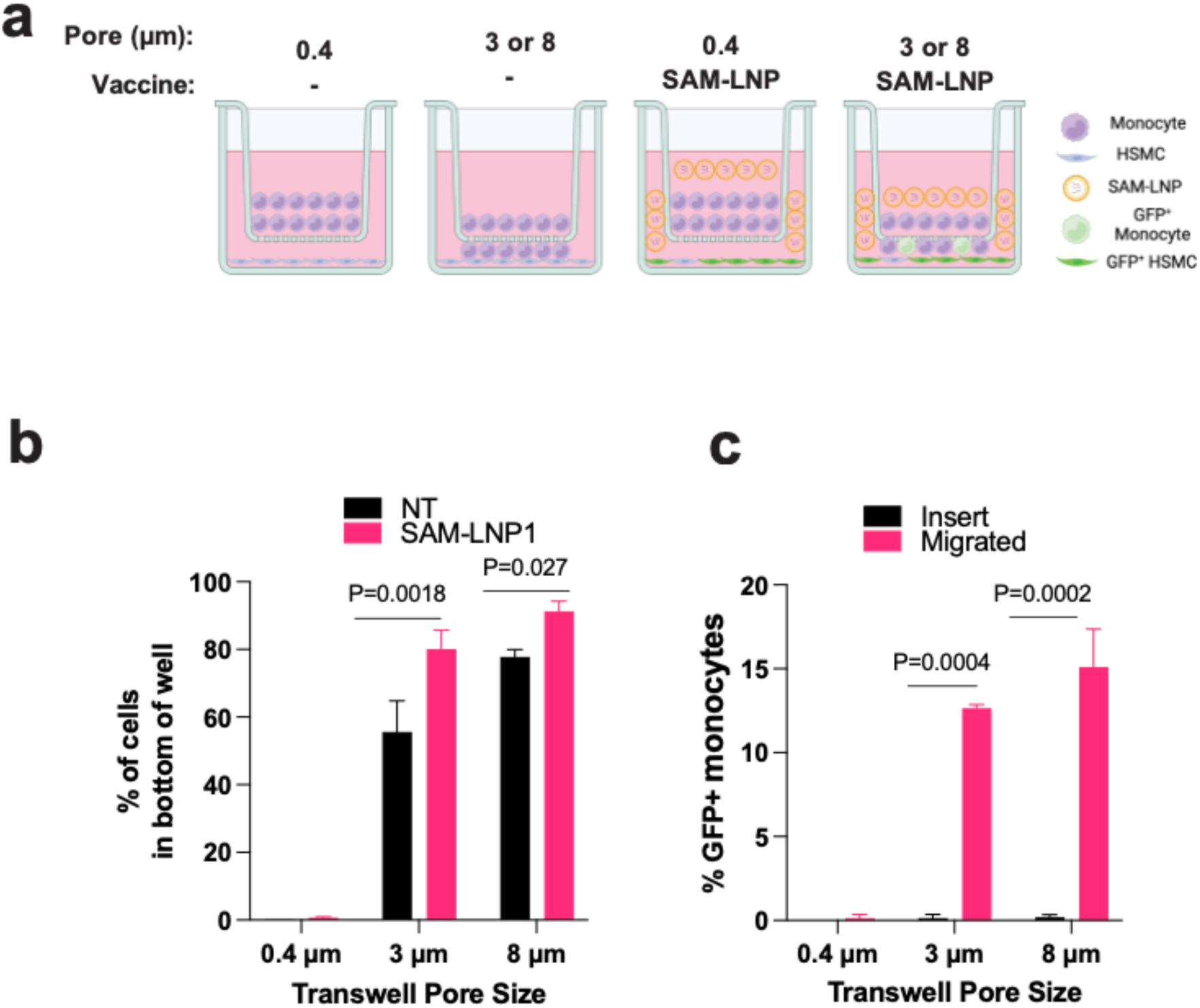
Requirement of direct cell-cell contact for antigen expression induced by SAM-LNP1. a, Schematic of the TW experiment designed to test antigen uptake by monocytes and HSMC. **b,** Total monocytes recovered from both the insert and the bottom of TWs with 0.4 µm, 3 µm, and 8 µm pore sizes with no treatment (NT) or 100 ng/ml SAM-LNP1, collected separately and quantified via flow cytometry. **c,** Percentage of GFP+ monocytes in the insert or those that migrated to the HSMC monolayer in the bottom wells relative to the total number of monocytes collected from each location. Results from one donor are shown; similar outcomes were achieved in two additional experiments using monocytes from two different donors. Data shown are mean ± SD; n = 2 replicates in each group in **b** and **c**, unpaired t-test.

**Supplementary Fig. 4.**
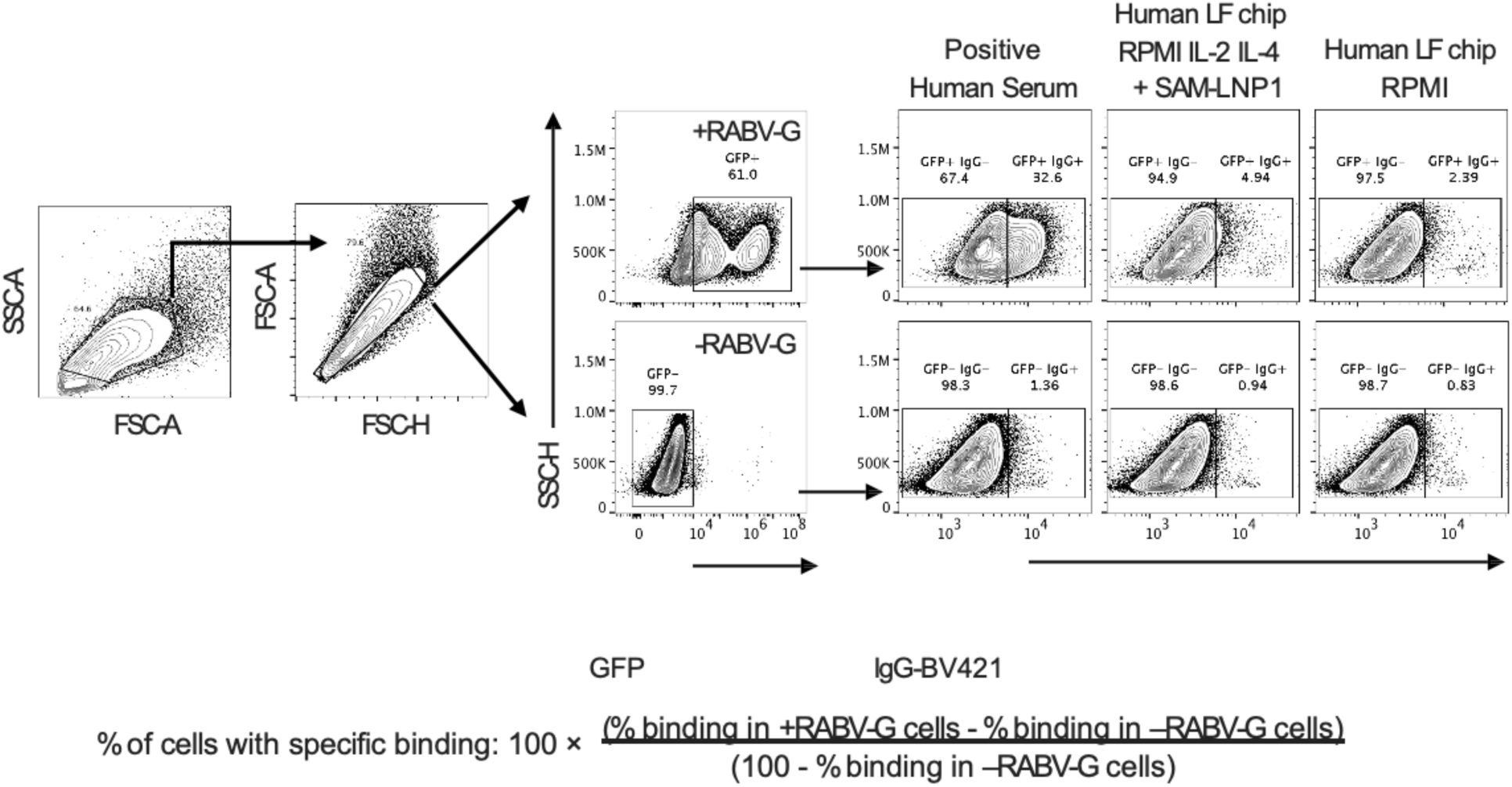
Detection and quantification of anti-RABV-G IgG using a cell-based assay. Gating strategy used for quantifying anti-RABV-G IgG, including the formula employed to determine IgG levels.

**Supplementary Fig. 5.**
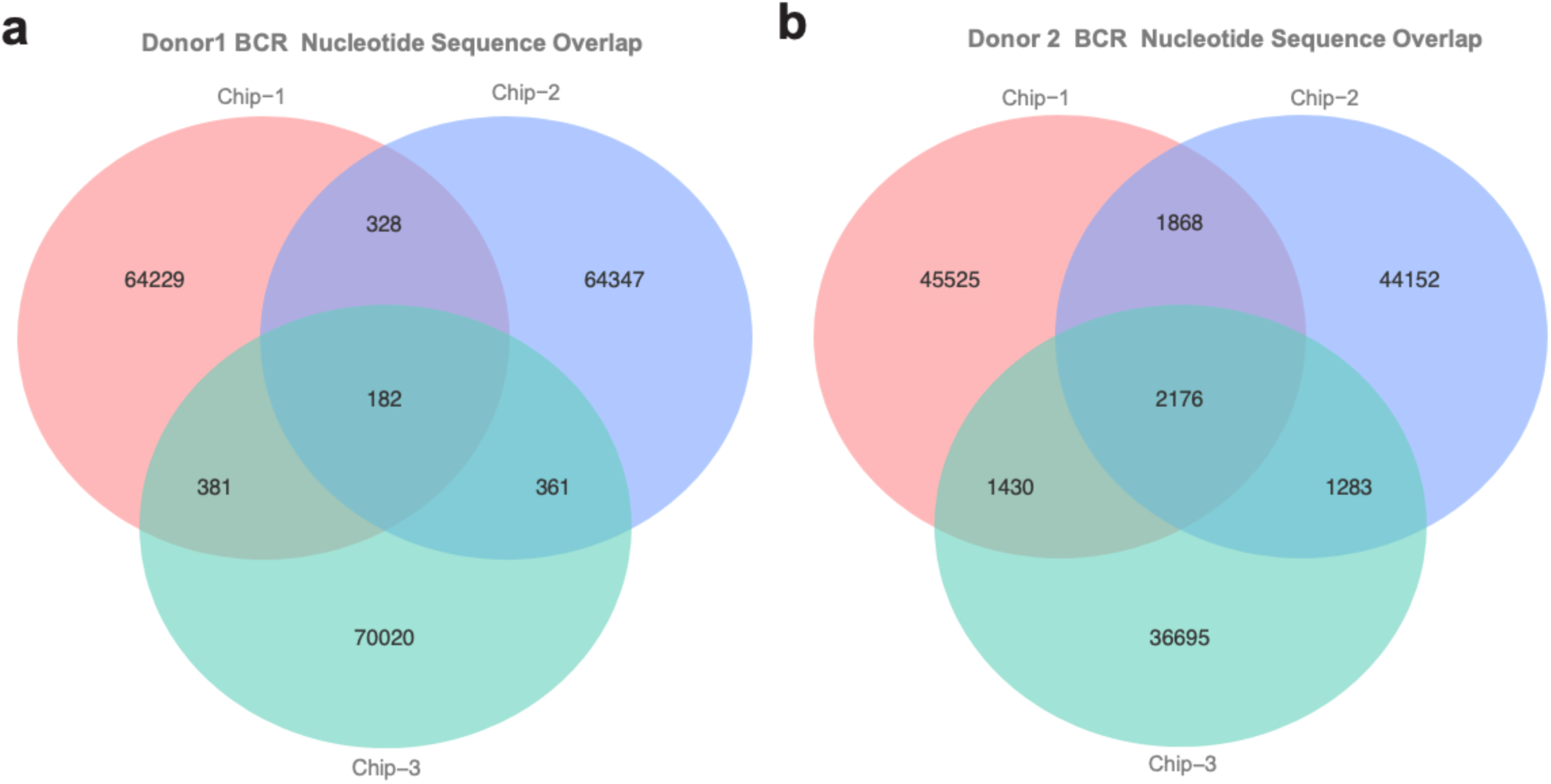
Overlap of BCR nucleotide sequences in vaccinated LF chips across two donors. Analysis of BCR sequence commonality in LF chips from individual donors following Moderna bivalent COVID-19 vaccine administration. **a,** BCR nucleotide sequence overlap in three LF chips from donor 1. **b,** BCR nucleotide sequence overlap in three LF chips from donor 2.

**Supplementary Fig. 6.**
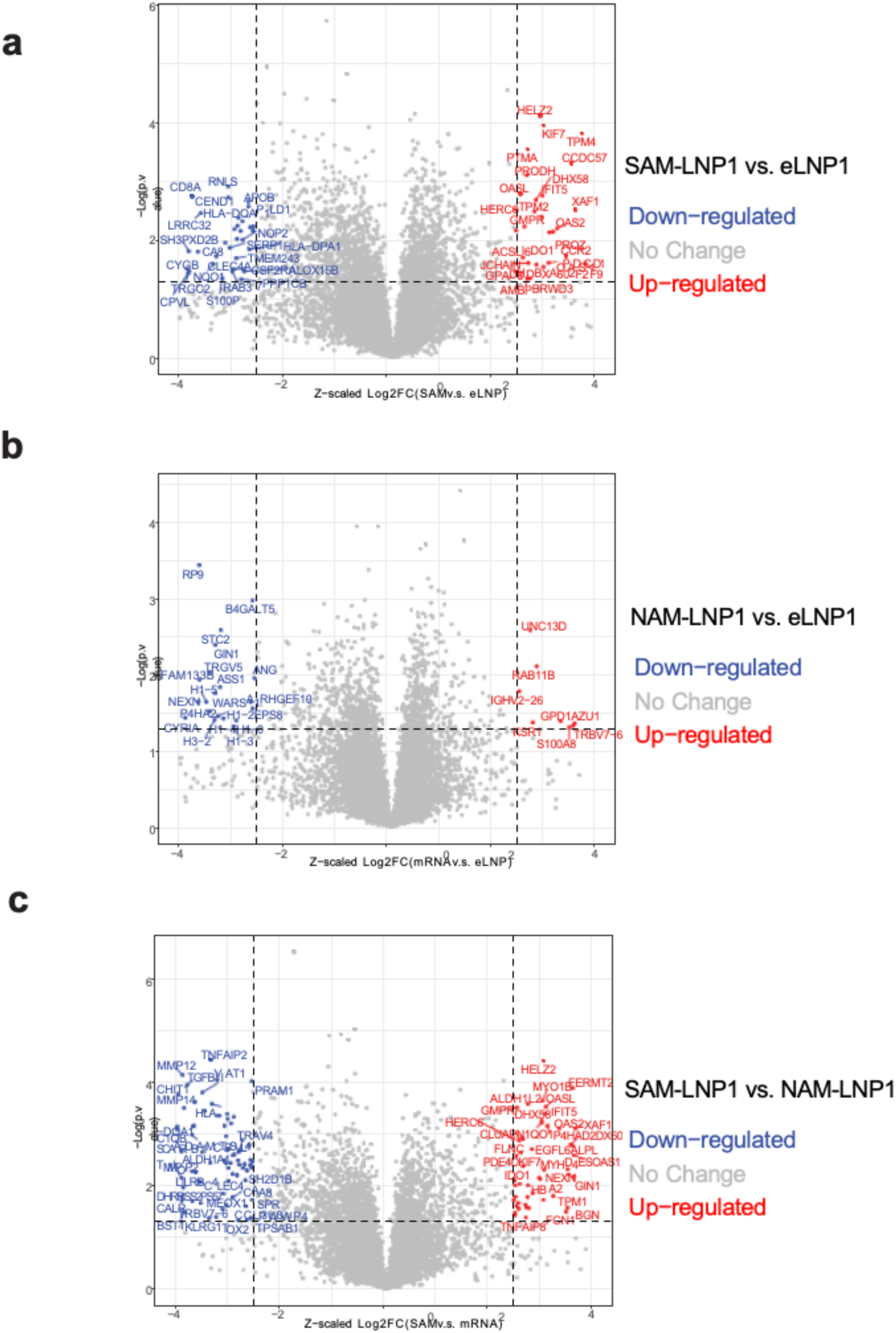
Comparative analysis of proteasome changes in vaccinated chips. Differential protein expression was assessed in LF chips following vaccination and visualized through volcano plots. **a,** Volcano plot illustrating protein changes in cells from chips vaccinated with SAM-LNP1 versus those treated with control empty LNP. **b,** Volcano plot showing protein expression in cells from chips vaccinated with NAM-LNP1 compared to control empty LNP treated chips. **c,** Volcano plot comparing protein expression in cells from SAM-LNP1 vaccinated chips to those vaccinated with NAM-LNP1. P-values were adjusted for multiple comparisons using the Bonferroni correction.

